# Accelerated Cortical Thinning within Structural Brain Networks is Associated with Irritability in Youth

**DOI:** 10.1101/596346

**Authors:** Robert J. Jirsaraie, Antonia N. Kaczkurkin, Sage Rush, Kayla Piiwia, Azeez Adebimpe, Danielle S. Bassett, Josiane Bourque, Monica E. Calkins, Matthew Cieslak, Rastko Ciric, Philip A. Cook, Diego Davila, Mark A. Elliott, Ellen Leibenluft, Kristin Murtha, David R. Roalf, Adon F.G. Rosen, Kosha Ruparel, Russell T. Shinohara, Aristeidis Sotiras, Daniel H. Wolf, Christos Davatzikos, Theodore D. Satterthwaite

**Affiliations:** Department of Psychiatry, Perelman School of Medicine, University of Pennsylvania, Philadelphia, PA 19104, USA; Department of Bioengineering, School of Engineering and Applied Science, University of Pennsylvania, Philadelphia, PA 19104, USA; Department of Electrical & Systems Engineering, School of Engineering and Applied Science, University of Pennsylvania, Philadelphia, PA 19104, USA; Department of Physics & Astronomy, College of Arts & Sciences, University of Pennsylvania, Philadelphia, PA 19104, USA; Department of Neurology, Perelman School of Medicine, University of Pennsylvania Philadelphia, PA 19104, USA; Department of Radiology, Perelman School of Medicine, University of Pennsylvania, Philadelphia, PA 19104, USA; Section on Mood Dysregulation and Neuroscience, National Institute of Mental Health (NIMH), 9000 Rockville Pike, Bethesda, MD, 20892, USA; Department of Biostatistics, Epidemiology, and Informatics, University of Pennsylvania, Philadelphia, PA 19104, USA

## Abstract

**Background:** Irritability is an important dimension of psychopathology that spans multiple clinical diagnostic categories, yet its relationship to patterns of brain development remains sparsely explored. Here, we examined how trans-diagnostic symptoms of irritability relate to the development of structural brain networks.

**Methods:** All participants (n=144, 87 females) completed structural brain imaging with 3 Tesla MRI at two timepoints (mean age at follow-up: 20.9 years, mean inter-scan interval: 5.1 years). Irritability at follow-up was assessed using the Affective Reactivity Index, and cortical thickness was quantified using Advanced Normalization Tools software. Structural covariance networks were delineated using non-negative matrix factorization, a multivariate analysis technique. Both cross-sectional and longitudinal associations with irritability at follow-up were evaluated using generalized additive models with penalized splines. The False Discovery Rate (*q*<0.05) was used to correct for multiple comparisons.

**Results:** Cross-sectional analysis of follow-up data revealed that 11 of the 24 covariance networks were associated with irritability, with higher levels of irritability being associated with thinner cortex. Longitudinal analyses further revealed that accelerated cortical thinning within 9 networks was related to irritability at follow-up. Effects were particularly prominent in brain regions implicated in emotion regulation, including the orbitofrontal, lateral temporal, and medial temporal cortex.

**Conclusions:** Collectively, these findings suggest that irritability is associated with widespread cortical thickness reductions and accelerated cortical thinning, particularly within frontal and temporal cortex. Aberrant structural maturation of regions important for emotional regulation may in part underlie symptoms of irritability.

## INTRODUCTION

Irritability is a debilitating dimension of psychopathology that cuts across multiple psychiatric disorders, including major depression, bipolar disorder, generalized anxiety disorder, oppositional defiant disorder, and disruptive mood dysregulation disorder. Symptoms of irritability include angry mood, exaggerated responses to negative stimuli, and limited frustration tolerance [1]. While many youths with severe irritability are diagnosed with bipolar disorder, studies utilizing careful assessment and longitudinal follow-up have demonstrated that the phenotype of chronic non-episodic irritability is distinct from bipolar disorder [2–6]. This conceptualization is supported by evidence that symptoms of irritability exist along a continuum and can be measured reliably across disorders as a unique dimension [7–9]. Prior studies have revealed that irritable youths are at increased risk for anxiety and depressive disorders and diminished functional outcomes, including reduced income and educational attainment [2, 10, 11]. Despite growing research on the impact of irritability, its neurobiological substrates remain sparsely investigated. This gap is critical, as understanding how aberrant patterns of brain development confer vulnerability to irritability is a prerequisite for targeted interventions that “bend the curve” of brain maturation to achieve better outcomes.

Symptoms of irritability are common in early childhood but generally decrease with age [12]. However, there are subgroups of children who exhibit chronically elevated levels of irritability throughout development [12–14]. To date, only two studies have investigated how irritability relates to longitudinal brain development [15, 16]. Adleman et al. (2012) reported that irritable youths had decreased gray matter volume in regions that support cognitive control and motor inhibition [17], such as the pre-supplementary motor area, insula, and dorsolateral prefrontal cortex. In contrast, Pagliaccio et al. (2018) reported that chronically irritable youths had increased cortical thickness in the superior temporal and dorsolateral prefrontal cortex. Notably, longitudinal analyses from both studies did not detect any significant differences in the rate of cortical changes between the irritable group and typically developing groups. However, both studies used standard image analysis techniques and assessed irritability using a case-control design.

In addition to this prior work in structural imaging, studies using functional imaging have found that chronic irritability is associated with aberrant activation in fronto-temporal regions important for emotion regulation, including the dorsolateral prefrontal cortex, lateral orbitofrontal cortex, and anterior temporal cortex [18–20]. Additional functional MRI studies using response inhibition tasks have similarly reported that irritability is related to abnormal activation of the dorsolateral prefrontal cortex, which could further contribute to deficits in affect regulation [21, 22]. Studies using functional near infrared spectroscopy suggest that abnormalities of the dorsolateral prefrontal cortex can be detected as early as 3-5 years of age [23, 24]. Furthermore, anger and irritability were found to be inversely correlated with the functional connectivity between the amygdala and regulatory regions such as the orbitofrontal cortex [25–27].

Taken together, these findings suggest that abnormalities within executive and affective circuity may be associated with chronic irritability. However, longitudinal studies of irritability are scarce, thereby limiting inferences that directly relate irritability to development. In fact, previous longitudinal studies on irritable youths have only acquired neuroimaging data between the ages of 7 to 16 years [15, 16]. Consequently, less is known about the neural markers of irritability during both infancy and early adulthood. Both stages of development are important given that the fronto-temporal regions critical for affect regulation undergoes protracted maturation throughout childhood and well into early adulthood [9, 28]. In particular, mapping neural trajectories as irritable youths transition into adulthood could be helpful for preventing deleterious outcomes during that period, which is when substance use, mood, and psychotic disorders often emerge [29].

In this study, we investigated how longitudinal structural brain maturation relates to a dimensional measure of irritability in a trans-diagnostic sample of adolescents and young adults. We hypothesized that irritability at follow-up would be associated with reduced cortical thickness and accelerated longitudinal thinning in brain regions implicated in emotion regulation such as the dorsolateral, ventrolateral, and orbitofrontal cortex. In contrast to prior studies that examined structural differences within specific regions [15] or across hundreds of thousands of voxels [16], we studied structural covariance networks using a powerful multivariate analysis technique known as non-negative matrix factorization (NMF). Previous work using this approach has demonstrated that NMF-derived networks enhance both interpretability and statistical power [30, 31]. As described below, we provide new evidence that irritability is associated with reduced cortical thickness and accelerated thinning in brain regions critical for emotion regulation.

## METHODS

### Participants

A total of 144 participants (mean age = 15.9 years, SD = 3, range = 8-22 years, 82 non-Caucasian, 87 females) were recruited from the Philadelphia Neurodevelopment Cohort (PNC; [32, 33]). As part of the PNC, all participants received a psychiatric screening interview and a structural MRI scan. Four items from the psychiatric interview provided a baseline screening for irritability (see **Supplementary Methods** for details). Participants who screened positive for at least one symptom of irritability were more likely to be recruited for a follow-up visit, in order to obtain a sample enriched with irritability across both timepoints. Of the participants who had usable data at both timepoints, 102 endorsed at least one symptom of irritability and 35 did not screen positive for any symptoms at baseline (see Table 1). Follow-up diagnostic interviews, irritability assessments, and MRI scans were completed with a mean inter-scan interval of 5.1 years (inter-scan interval SD = 1.2, mean age = 20.9 years, SD = 3.0, range = 13-26 years). All procedures were approved by the Institutional Review Boards of the University of Pennsylvania and the Children’s Hospital of Philadelphia.

**Table 1.** Sample Demographics and Clinical Characteristics.

| Characteristics | Non-Irritable at Baseline<br>(n=35) | Irritable at Baseline<br>(n=102) | Statistics<br>(t, $X^2$) | p-value |
| --- | --- | --- | --- | --- |
| Age at Baseline (Years) | 15.4 (3.7) | 16.2 (3.0) | 1.12 | .266 |
| Age at Follow-up (Years) | 20.4 (3.5) | 21.3 (2.7) | 1.44 | .157 |
| Sex (% of Females) | 45.7 | 37.3 | 0.47 | .495 |
| Father Education (Years) | 15.0 (2.9) | 13.8 (2.6) | 2.12 | .039 |
| Mother Education (Years) | 15.3 (2.3) | 14.4 (2.6) | 2.01 | .048 |
| Irritability (ARI scores) | 2.0 (1.6) | 3.4 (2.9) | 3.70 | <.001 |
| Depression (BDI scores) | 2.6 (3.0) | 6.0 (7.6) | 3.68 | <.001 |
| ADHD (SWAN scores) | 1.7 (2.1) | 3.7 (3.9) | 3.75 | <.001 |
| Anxiety (SCARED scores) | 10.8 (8.1) | 15.5 (12.1) | 2.56 | .012 |
| Clinically Diagnosed at Baseline (%) | 22.9 | 88.2 | 51.5 | <.001 |
| Clinically Diagnosed at Follow-up (%) | 22.9 | 60.9 | 13.5 | <.001 |

*Note.* All variables are presented as *M* (SD), except for sex and diagnostic status at both timepoints, which are presented as the percentage of participants within a given group. For specific details regarding clinical diagnoses, please refer to Table S1.

### Clinical Phenotyping

Psychopathology symptoms at baseline were evaluated using a structured computerized screening interview [34], which was a modified version of the Kiddie-Schedule for Affective Disorders and Schizophrenia [35]. As previously described [36], the GOASSESS interview collected information on symptoms, frequency, duration, distress, and impairment for lifetime mood, anxiety, behavioral, eating, and psychosis spectrum disorders. A total of 39 participants did not screen positive for any psychiatric disorder, and 99 screened positive for at least one psychiatric disorder. Notably, there were not any specialized assessments of irritability collected baseline. However, the four items from the GOASSESS screening interview were summed to provide a coarse dimensional assessment of irritability at baseline (see **Supplementary Methods)**.

Clinical diagnoses at follow-up were assessed using a custom protocol [37] including modules from the Kiddie-Schedule for Affective Disorders and Schizophrenia [35], Disruptive Mood Dysregulation Disorder Child Version [20], and the psychotic and mood differential diagnosis modules of the Structured Clinical Interview for DSM-IV [38]. All sections were administered by highly-trained assessors in a semi-structured manner, allowing for follow-up probes and clarification of endorsed items. At the second timepoint, there were 67 participants who did not meet criteria for a diagnosis and 70 participants with at least one clinical diagnosis (see Table S1).

At follow up, the Affective Reactivity Index (ARI) was used to measure self-reported symptoms of irritability [39]. This scale contains six items on symptom severity and a seventh item on functional impairment, which are rated on a three-point scale (ranging from 0 for “not true” to 2 for “certainly true”). The reliability and validity of the ARI have been previously reported [39, 40]. The sum of the first 6 items on the self-report was used as a dimensional measure of irritability, which was log transformed to obtain a more normal distribution.

As psychiatric symptoms have substantial comorbidity, one potential risk is that any observed associations with irritability may be driven by collinearity with other symptom domains. To evaluate this possibility, symptoms from relevant domains were measured at follow-up using the Beck Depression Inventory (BDI; [41]), Screen for Child Anxiety Related Disorders (SCARED; [42]) and Strengths and Weaknesses of ADHD-symptoms and Normal-behaviors (SWAN; [43]). These additional dimensions of psychopathology were included as covariates in sensitivity analyses.

### Image acquisition

The baseline structural images were collected as part of the Philadelphia Neurodevelopment Cohort [32, 33]. In brief, all baseline scans were acquired on the same MRI scanner (Siemens TIM Trio 3 Tesla, Erlangen, Germany; 32-channel head coil) using the same imaging sequence. Structural brain imaging was completed using a magnetization-prepared, rapid acquisition gradient-echo (MPRAGE) T1-weighted image with the following parameters: TR 1810 ms; TE 3.51 ms; FOV 180×240 mm; matrix 256×192; 160 slices; slice thickness/gap 1/0 mm; TI 1100 ms; flip angle 9 degrees; effective voxel resolution of 0.93 × 0.93 × 1.00 mm; total acquisition time 3:28 min. The follow-up structural scans were completed on a different MRI scanner (Siemens Prisma 3 Tesla, Erlangen, Germany; 64-channel head coil), but the imaging sequence and parameters were identical to those used on the baseline acquisition, and all measures were statistically harmonized prior to analysis; we describe this harmonization procedure in more detail below.

### Image processing and Quality Assurance

Structural image processing for estimating cortical thickness (CT) was completed with Advanced Normalization Tools version 2.1.0 (ANTs). A custom template and tissue priors were utilized to avoid registration bias and maximize sensitivity to detect regional effects that can be impacted by registration errors. Structural images were then processed and registered to this template using the ANTs CT pipeline [44, 45]. This procedure includes brain extraction, N4 bias field correction [46], Atropos probabilistic tissue segmentation [47], top-performing SyN diffeomorphic registration to template space [48], and diffeomorphic registration based cortical thickness measurement [49]. The ANTs CT pipeline has been validated in multiple open data sets and found to have good test-retest reproducibility and enhanced ability to predict age and gender from regional thickness measurements when compared to FreeSurfer [44]. CT images were down-sampled to 2 mm voxels and subsequently smoothed by 4 mm before applying non-negative matrix factorization.

All raw and processed images were reviewed by highly trained image analysts who identified unusable images at baseline (n=4) and follow up (n=3) as well as provided a quantitative quality score for each image [50]. Every analyst was trained to >85% concordance with faculty consensus ratings on an independent dataset. Images with substantial artifact were excluded from analyses, resulting in a baseline sample of 140 participants, a follow-up sample of 141 participants, and a longitudinal sample of 137 participants.

### Non-negative matrix factorization

Cortical thickness data from both timepoints were summarized using structural covariance networks for two reasons. First, prior work has shown inherent patterns of covariance in brain structure and analyzing the data according to this covariance structure enhances interpretability [51, 52]. Second, distilling the data into covariance networks reduces the large number of multiple comparisons present in voxel-based morphometry studies, and therefore reduces both Type I errors (if insufficient correction is applied) and Type II errors (if conservative correction is applied) [53]. We used non-negative matrix factorization (NMF) to identify structural networks in which CT consistently co-varies across individuals and brain regions (see Figure 1A). This tool (https://github.com/asotiras/brainparts) implements an extension of standard NMF that adopts orthonormality constraints for the estimated structural covariance networks, and projective constraints for their respective loadings [54]. This procedure yields compact networks with positive weights that are more interpretable and reproducible compared to other decomposition techniques, such as principal component analysis and independent component analysis [30, 31]. Additionally, this procedure employs a deterministic initialization algorithm [55] that leads to faster convergence, promotes sparsity, and ensures consistent results across multiple runs.

**Figure 1.**
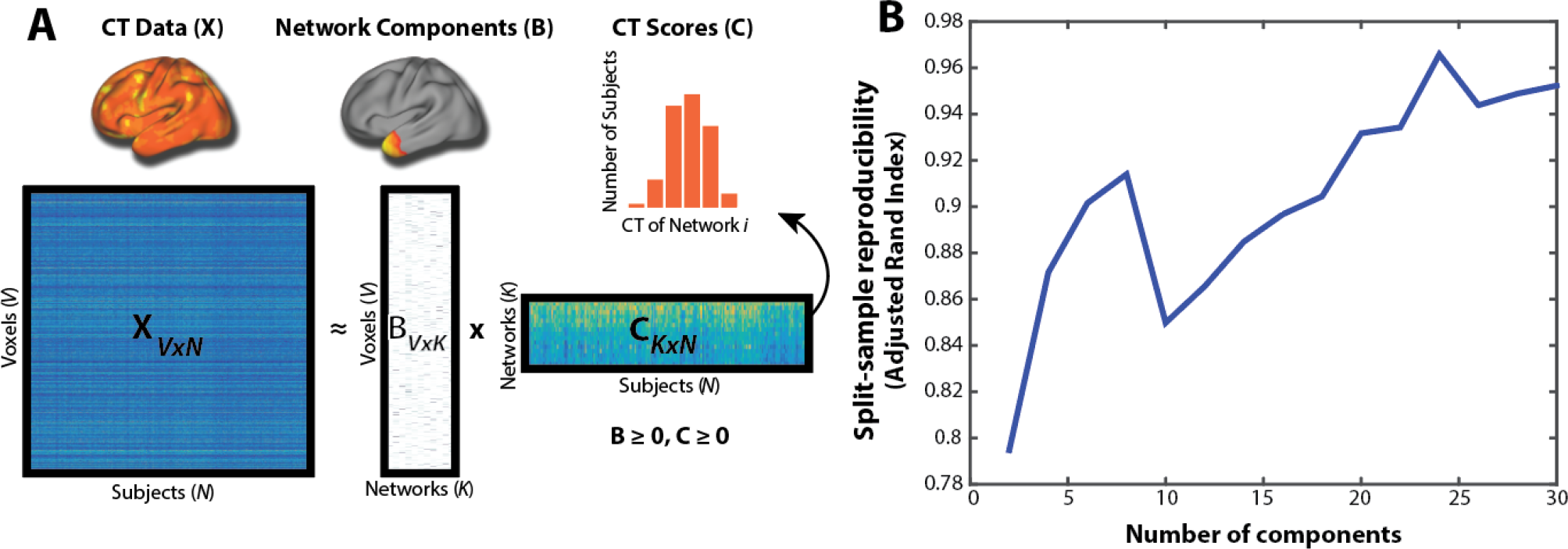
Non-Negative Matrix Factorization. **A**. Schematic of non-negative matrix procedure. The *X* matrix represents the cortical thickness (CT) data (columns) for all subjects (rows). The *B* matrix represents estimated networks (columns) and their leadings on each voxel (rows). The *C* matrix provides the subject-specific weights (columns) for each network (rows); the histogram shows CT scores in a single network that corresponds to a row in the *C* matrix. Matrix sizes are shown with following dimensions: *V* = number of CT voxels, *N* = number of participants; *K* = number of networks. **B**. The 24-network resolution was selected on this basis of split-half reproducibility, as measured by the Adjusted Rand Index.

Multiple NMF resolutions were examined (2 to 30 networks, in increments of 2) in order to select the ideal number of components. Each resolution was computed in a split-half sample (matched on age, sex, image quality, and irritability) to evaluate reproducibility. The optimal number of components (24) was chosen based on the solution with the highest reproducibility (96.5%), which was quantified using the Adjusted Rand Index (see Figure 1B). NMF networks are three-dimensional but were mapped to a surface for visualization using Caret (see Figure 2; [56]).

**Figure 2.**
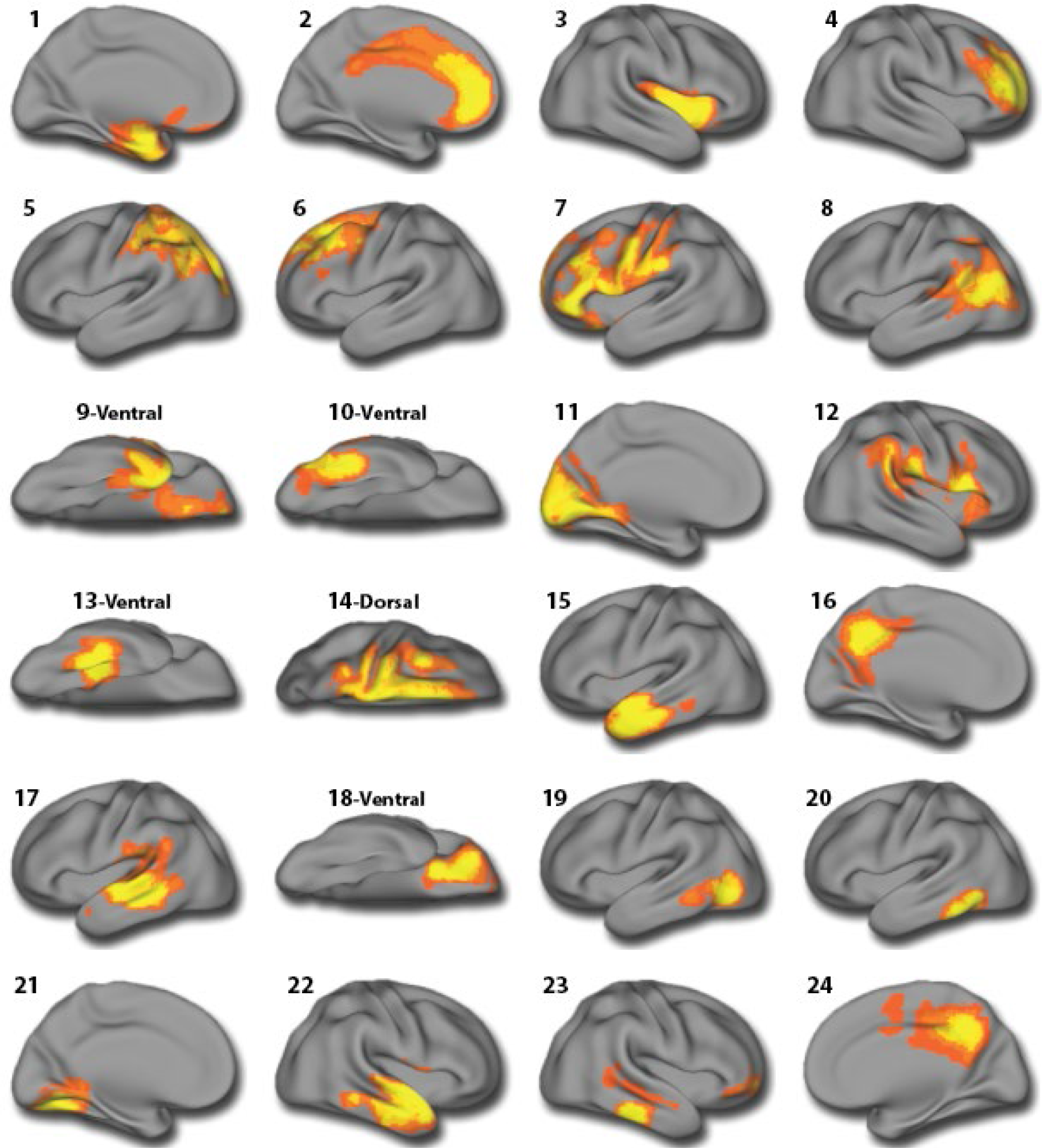
Structural covariance networks. Visualization of the 24 structural covariance networks revealed by NMF. Warmer colors indicate higher loadings. Networks include 1) Medial temporal lobe, 2) Anterior cingulate, 3) Insula, 4) Anterior prefrontal cortex, 5) Posterior parietal cortex, 6) Dorsolateral prefrontal cortex, 7) Inferior frontal cortex, 8) Temporo-parietal junction, Temporal pole and medial orbitofrontal cortex, 10) Inferior temporal gyrus, 11) Occipital cortex, 12) Peri-sylvian cortex, 13) Fusiform gyrus, 14) Dorsal frontoparietal cortex, 15) Lateral temporal cortex, 16) Posterior cingulate cortex, 17) Supramarginal gyrus, 18) Lateral orbitofrontal cortex, 19) Temporo-occipital cortex, 20) Inferior temporal cortex, 21) Lingual gyrus, 22) Superior temporal gyrus, 23) Inferior temporal sulcus, 24) Precuneus.

### Statistical Harmonization

To minimize potential variance introduced by different MRI scanners, the cortical thickness values of structural brain networks from both timepoints were harmonized using the ComBat procedure, which capitalizes upon Bayesian techniques that were originally implemented in the context of statistical genomics [57]. Subsequent studies using neuroimaging data have shown that ComBat removes unwanted sources of scanner variability and preserves biological associations in the data [58–60]. Effects of age, sex, and irritability were protected in the harmonization process.

### Statistical analyses

As described in detail below, we evaluated cross-sectional relationships between irritability and CT at both timepoints, in addition to the longitudinal associations between irritability at follow-up and the annualized rates of change in CT. We focused on associations with irritability at follow-up since our assessment of irritability at baseline was limited (four dichotomous screening items). Statistical analyses were completed using R version 3.5.1 [61], and all code is publicly available (https://github.com/PennBBL/jirsaraieStructuralIrritability). Given that brain development is a non-linear process [29, 62], in all analyses age was modeled using penalized splines within a generalized additive model [63]. In this type of model, a penalty is assessed on nonlinearity using restricted maximum likelihood in order to avoid overfitting. The false discovery rate was controlled (*q*<0.05) using the Benjamini-Hochberg procedure to correct for multiple comparisons.

As a first step, we examined the relationships at baseline between CT and the sum of the four irritability screening items. Second, we evaluated the relationships at follow-up between CT and irritability as measured by the ARI. Third, we evaluated how irritability at follow-up (using the ARI) related to longitudinal rates of change. Cross-sectional analyses included age, sex, and quality assurance ratings as covariates; longitudinal analyses included age at baseline, sex, and an average of the quality assurance ratings across timepoints as covariates. Longitudinal analyses evaluated the annualized rates of change in CT to account for varying inter-scan intervals [64]. The total difference in CT between both timepoints was divided by baseline CT to get the total percent change (Δ*CT*⁄*CT*_*TP*1_), which was subsequently divided by the time between scans(Δ*Age*), yielding standardized the percent change of CT across all participants(*CT*_*Rate*_):

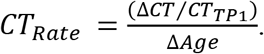

Sensitivity analyses were conducted on all significant results to ensure that effects were not being driven by maternal education, diagnosis status, or other dimensions of psychopathology (see above). Each of these variables were included as model covariates. Furthermore, two additional analyses were conducted excluding participants taking psychotropic medications or using recreational drugs. Finally, we examined interactions between irritability and diagnostic status (typically developing vs. those with any psychiatric disorder), irritability and sex, as well as irritability and age.

## RESULTS

### Irritability is elevated across diverse diagnostic categories

Irritability at follow-up was not associated with demographic variables at follow-up, such as age (*p*=.657), sex (*p*=.431), participant education (*p*=.490), paternal education (*p*=.166), or maternal education (*p*=.872). As expected, irritability at follow-up varied among diagnostic groups at both baseline (*p*=.028) and follow-up (*p*<.001). However, this effect was completely driven by differences between the typically developing participants and those who met diagnostic criteria for a psychiatric disorder; irritability did not differ between specific psychiatric disorders at baseline (*p*=.535) or follow-up (*p*=.446).

### Cross-sectional irritability at follow-up is associated with thinner cortex

As an initial step, we assessed cross-sectional associations between irritability and cortical thickness at each timepoint. At baseline, we examined the limited measure of irritability that was available (sum of four screening items), whereas the cross-sectional analysis in follow-up data used the specialized assessment provide by the ARI. Cross-sectional analyses of this coarse baseline data did not reveal any associations that survived FDR-correction. In contrast, dimensional irritability at follow-up was associated with thinner cortex within 11 of the 24 NMF networks (see Table 2). Effects were particularly prominent in networks that included the orbitofrontal cortex, which is critically involved in emotion regulation (see Figure 3). Moderation analyses revealed that the relationship between irritability at follow-up and the dorsolateral prefrontal cortex (network 6) weakened with age (see Figure S1); no other interactions were significant.

**Table 2.** Cross-sectional associations between irritability at follow-up and structural covariance networks at follow-up.

| NMF-Networks | $\beta$ | *SE* | *t* | *p* | *p-FDR* |
| --- | --- | --- | --- | --- | --- |
| N-1: Medial temporal lobe | -0.28 | 1.17 | -3.79 | <.001 | .002 |
| N-8: Temporo-parietal junction | -0.22 | 1.44 | -2.94 | .004 | .011 |
| N-9: Temporal pole & medial orbitofrontal cortex | -0.20 | 1.30 | -2.71 | .008 | .023 |
| N-10: Inferior temporal gyrus | -0.21 | 1.34 | -2.66 | .009 | .029 |
| N-11: Occipital cortex | -0.26 | 1.71 | -3.40 | <.001 | .005 |
| N-13: Fusiform gyrus | -0.35 | 1.07 | -4.74 | <.001 | <.001 |
| N-15: Lateral temporal cortex | -0.31 | 1.10 | -4.10 | <.001 | <.001 |
| N-18: Lateral orbitofrontal cortex | -0.26 | 0.84 | -3.46 | <.001 | .004 |
| N-20: Inferior temporal cortex | -0.18 | 1.01 | -2.32 | .022 | .049 |
| N-21: Lingual gyrus | -0.20 | 1.17 | -2.56 | .012 | .027 |
| N-22: Superior temporal gyrus | -0.22 | 0.98 | -2.93 | .004 | .013 |

*Note.* $\beta$ (standardized regression coefficients), standard errors, t-statistics, uncorrected and FDR-corrected p-values for significant main effects that irritability at follow-up had on CT at follow-up. All analyses were conducted contained 136 degrees of freedom.

**Figure 3.**
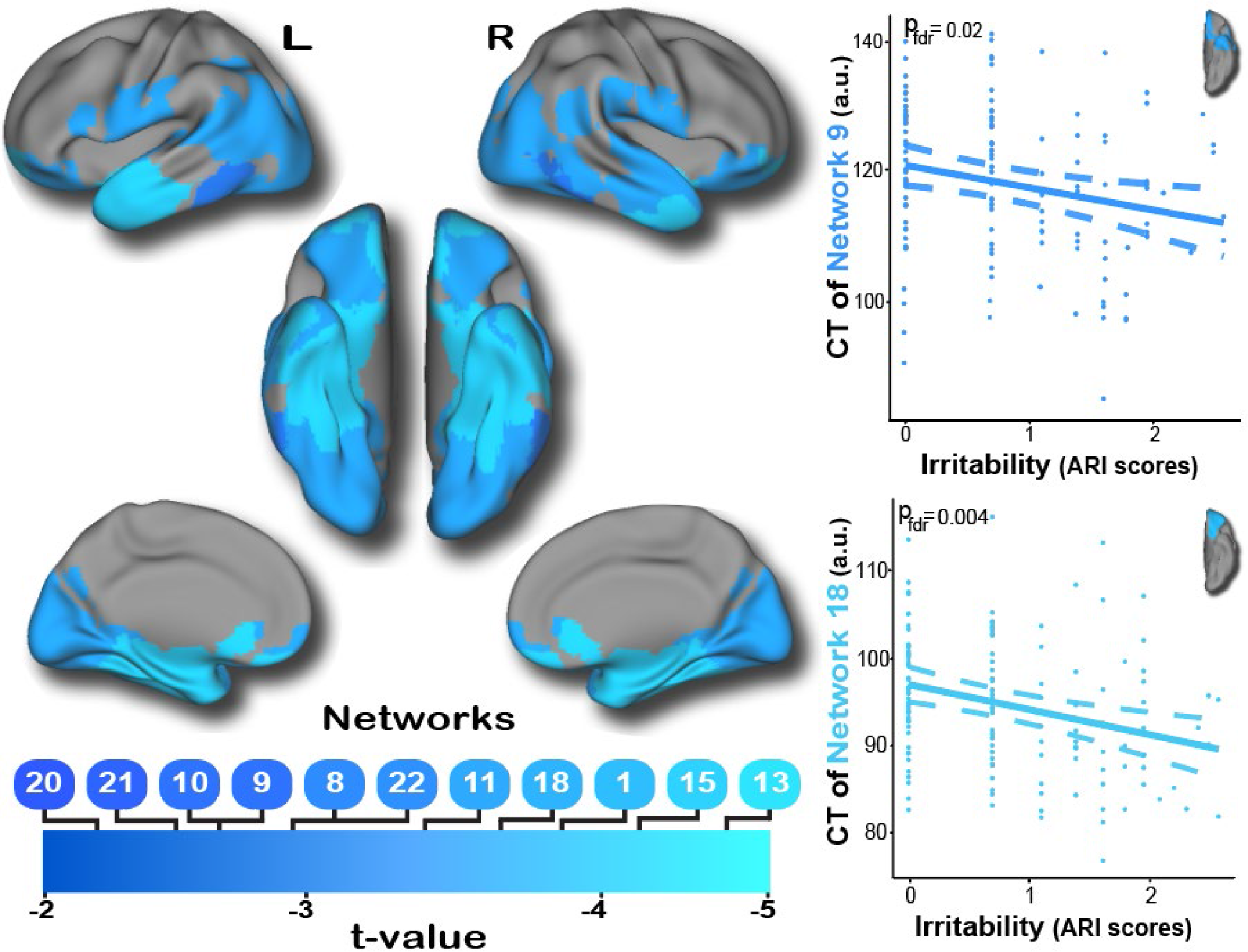
Thinner cortex within multiple structural covariance networks is associated with irritability at follow-up. Cross-sectional analyses reveal that thinner cortex within multiple frontal and temporal networks is associated with irritability at follow-up. These networks include regions critical for affective regulation, such as medial temporal cortex, temporal pole, and both medial and lateral orbitofrontal cortex.

### Sensitivity analyses provide convergent results

Sensitivity analyses of cross-sectional findings at follow-up revealed that irritability remained associated with all 11 networks across when controlling for maternal education (*P*_*FDR*_<.021), paternal education (*P*_*FDR*_<.032), or diagnostic status (*P*_*FDR*_<.049). Despite reduced statistical power, sensitivity analyses excluding 23 participants who were taking psychotropic medications did not significantly impact results, nor did excluding 32 participants who tested positive on a urine drug screen (primarily cannabis). Additional sensitivity analyses were conducted to determine whether these significant effects were specific to irritability or could reflect comorbid dimensions of psychopathology. Inclusion of depression or anxiety as covariates did not impact the significance of results; when dimensional ADHD symptoms were included as a covariate, 5 of the original 11 networks remained significant (see Table S2).

### Irritability at follow-up is associated with accelerated longitudinal cortical thinning

Finally, we evaluated associations between irritability at follow-up and the longitudinal rates of change within CT networks. These analyses revealed that irritability at follow-up was significantly associated with accelerated longitudinal cortical thinning within 9 networks (see Table 3). Notably, 7 of these networks overlapped with the cross-sectional associations at follow-up (see Figure 4). Sensitivity analyses revealed that irritability remained associated with rate of change within all 9 CT networks when controlling for maternal education (*P*_*FDR*_<.029), paternal education (*P*_*FDR*_<.022), and diagnostic status (except for network 18; *P*_*FDR*_<.058). All significant effects remained when excluding participants who were taking psychotropic medications (*P*_*FDR*_<.003); 4 networks persisted after excluding participants who used had a positive urine drug screen (see Table S3). Finally, irritability remained associated with CT in these networks when controlling for comorbid symptoms of ADHD (*P*_*FDR*_<.032); 4 networks remained significant when covarying for anxiety or depression (see Table S3). Age, sex, and diagnostic status did not moderate the relationship between irritability and network CT.

**Table 3.** Longitudinal associations between irritability at follow-up and annualized rates of change in structural covariance networks.

| **NMF-Networks** | **$\beta$** | ***SE*** | ***t*** | ***p*** | ***p-FDR*** |
| --- | --- | --- | --- | --- | --- |
| N-1: Medial temporal lobe | -0.36 | <0.01 | -4.46 | <.001 | <.001 |
| N-7: Inferior frontal cortex | -0.21 | <0.01 | -2.48 | .014 | .043 |
| N-9: Temporal pole and medial orbitofrontal cortex | 0.28 | <0.01 | -3.36 | .001 | .008 |
| N-11: Occipital cortex | -0.34 | <0.01 | 4.32 | <.001 | <.001 |
| N-15: Lateral temporal cortex | -0.23 | <0.01 | -2.70 | .008 | .038 |
| N-17: Supramarginal gyrus | -0.21 | <0.01 | -2.53 | .013 | .043 |
| N-18: Lateral orbitofrontal cortex | -0.20 | <0.01 | -2.45 | .016 | .043 |
| N-20: Inferior temporal cortex | -0.23 | <0.01 | -2.81 | .006 | .034 |
| N-22: Superior temporal gyrus | -0.21 | <0.01 | -2.44 | .016 | .043 |

*Note.* $\beta$ (standardized regression coefficient), standard errors, t-statistics, uncorrected and FDR-corrected p-values for main effects of irritability at follow-up on the annual rates of change within CT networks. All analyses were conducted contained 132 degrees of freedom

**Figure 4.**
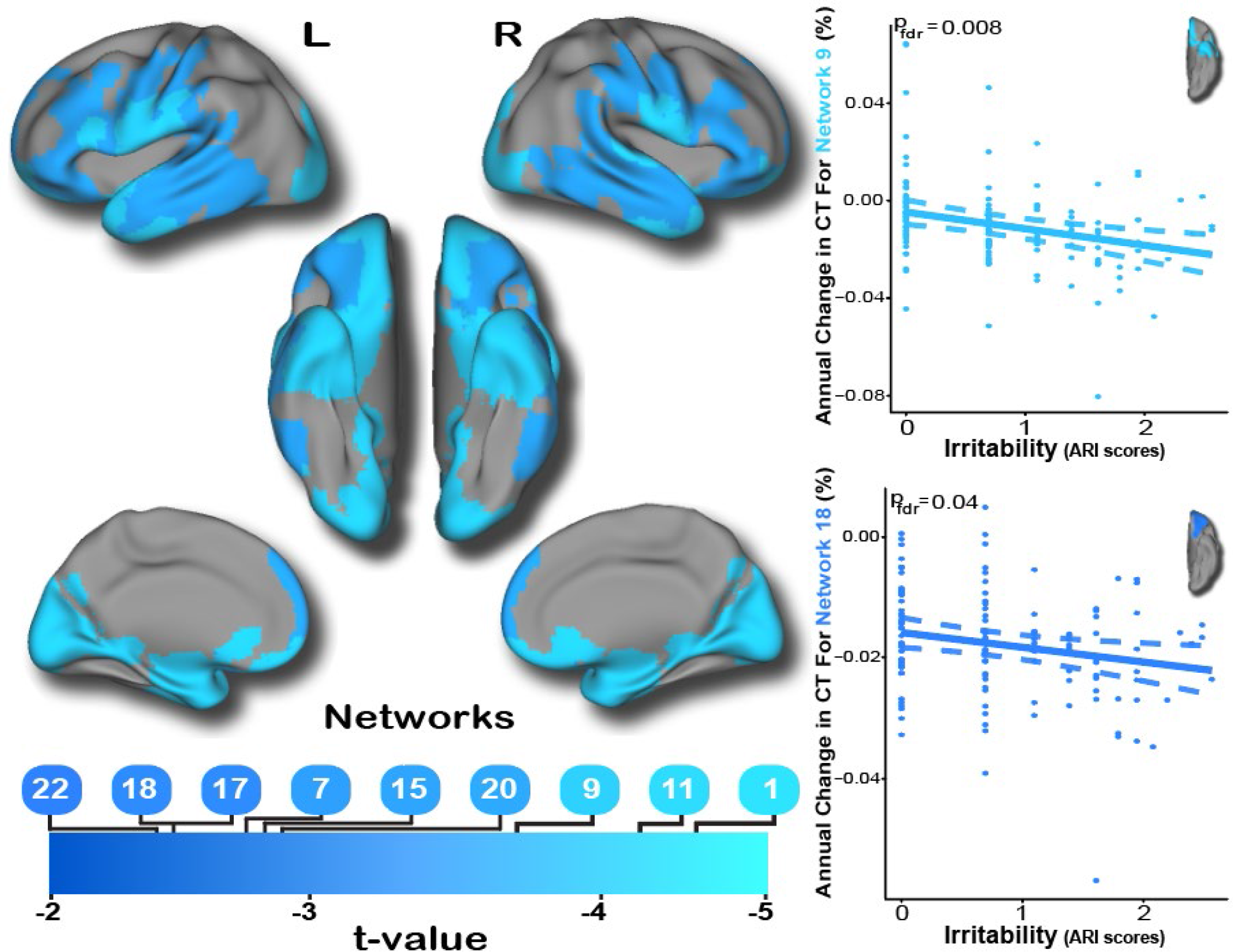
Accelerated cortical thinning is associated with irritability. Longitudinal analyses revealed that the annual rate of change in many frontal and temporal networks was also associated with irritability at follow-up. As for the cross-sectional analysis, these included the temporal pole, medial temporal cortex, and both medial and lateral orbitofrontal cortex.

## DISCUSSION

We evaluated the dimensional association between irritability in youth and cortical thickness within structural covariance networks. Relative to previous studies on the neurodevelopment of irritability [15, 16], we observed widespread effects across the frontal and temporal cortex. Of the 24 covariance networks derived from NMF, irritability was associated with thinner cortex within 11 networks at follow-up and accelerated longitudinal thinning within 9 networks. In general, these relationships did not vary by age, sex, or diagnosis status, and most relationships could not be explained by confounding variables or comorbid dimensions of psychopathology. The strongest main effects were observed in regions that are critical for emotion regulation, including orbitofrontal, lateral temporal, and medial temporal cortex. As described below, these results provide important insights regarding the neurodevelopmental substrates of trans-diagnostic irritability in youth.

Previous functional MRI studies have found that irritability is associated with aberrations in multiple brain regions, including the orbitofrontal cortex, medial temporal cortex, temporal pole, superior temporal gyrus, and occipital cortex [18–20]. We build upon this literature regarding brain function and demonstrate that many of these same regions exhibited cortical thickness reductions and accelerated cortical thinning that was related to trans-diagnostic symptoms of irritability. Structural deficits in regions necessary for affect regulation such as the orbitofrontal cortex and medial temporal cortex may be critical for the pathogenesis of irritability. Additionally, abnormalities in the fusiform gyrus and visual cortex may relate to deficits in facial emotion recognition, which has been robustly associated with irritability in prior work [20, 65–68]. Taken together, these findings further support the notion that deficits in inter-related brain networks underlie vulnerability to irritable mood.

Previous structural and functional MRI studies have suggested that irritability is related to abnormalities within the dorsolateral prefrontal cortex [15, 16, 18, 19, 23, 24]. Surprisingly, we did not find this association. However, previous studies were conducted on participants between the ages of 7 and 18, whereas our age range extended up to 26 years. Further, Tseng et al. (2018) reported that development was associated with a weakening of the relationship irritability and activation of the medial prefrontal cortex and anterior cingulate cortex during a frustration attention task. Thus, it is possible that the relationship between irritability and prefrontal cortex does not extend into early adulthood. Moderation analyses documenting an irritability-by-age interaction supported this possibility: among the younger participants in our sample, irritability was negatively associated with cortical thickness of the dorsolateral prefrontal cortex. However, this relationship waned by early adulthood.

The current study documented more robust and widespread associations between irritability and cortical thickness than previous structural MRI studies [15, 16, 69]. This difference may be attributed to three distinct advantages of the current approach. First, the structural covariance networks defined by NMF provided a parsimonious summary of the high-dimensional imaging data that limited multiple comparisons. This concise summary of the data allowed us to use rigorous FDR-correction for all comparisons, rather than cluster-based inferences that may be prone to Type 1 errors in many common implementations [70]. Second, instead of a categorical case-control approach, we used a validated dimensional measure of irritability. While case-control studies are useful for describing DMDD as a distinct clinical syndrome [1], a dimensional measurement of irritability may provide a greater sensitivity to detect individual differences in brain structure [9]. Lastly, most studies have analyzed irritability within a specific psychiatric disorder, such as bipolar disorder [16] or depression [15]. In comparison, relatively fewer studies have examined irritability in the context of the comorbidity that is commonly observed in clinical practice [19, 20, 68, 69]. Our study included a trans-diagnostic sample with substantial comorbidity, resulting in findings that may be more generalizable to the community.

Despite these advantages, several limitations should be noted. One disadvantage to having a sample with diverse clinical phenotypes and comorbid disorders is that we did not have sufficient statistical power to test whether the neural mechanisms underlying irritability differed among specific psychiatric disorders. Identifying the common and dissociable neural correlates of irritability across specific disorders could potentially have important implications for targeted therapies [1, 9]. Additionally, this study would have benefited from having the same dimensional measure of irritability at baseline and follow-up so that longitudinal analyses could have tested within-subject changes in irritability and CT. Finally, as previously noted, this is the first longitudinal study to examine brain maturation as irritable youths transition into adulthood. Ideally, longitudinal studies with three or more timepoints would map trajectories of brain development from childhood to adulthood.

In summary, we found that transdiagnostic symptoms of irritability were associated with widespread reductions in cortical thickness and accelerated cortical thinning in multiple brain networks. In particular, structural deficits were found in networks that support emotion regulation, including the orbitofrontal, medial temporal, and lateral temporal cortex. Future studies should incorporate repeated measurements of irritability and incorporate data from multi-modal imaging. Ultimately, such findings could be the basis for mapping the developmental substrates of irritability and allow for targeted interventions in youth at risk.

## SUPPLEMENTARY METHODS

The 4 GOASSESS items that were utilized as a baseline screening of irritability were from the depression, bipolar disorder, and oppositional defiant disorder sections of the interview. Each item is rated on a binary scale:

1. Has there ever been a time when you felt grouchy, irritable or in a bad mood most of the time; even little things would make you mad?
2. Has there ever been a time when you felt unusually grouchy, cranky, or irritable; when the smallest things would make you really mad?
3. Was there a time when you often did things that got you into trouble with adults such as losing your temper, arguing with or talking back to adults, or being grouchy or irritable with them?
4. Were you often irritable or grouchy, or did you often get angry because you thought that things were unfair?

The urine drug screen tested for multiple substances, including methamphetamines, opiates, PCP, benzodiazepines, barbiturates, amphetamines, cocaine, and cannabis. All 32 participants with a positive urine drug test screened positive for cannabis (THC) and 3 of these participants screened positive for another substance in addition to cannabis.

**Table S1.**
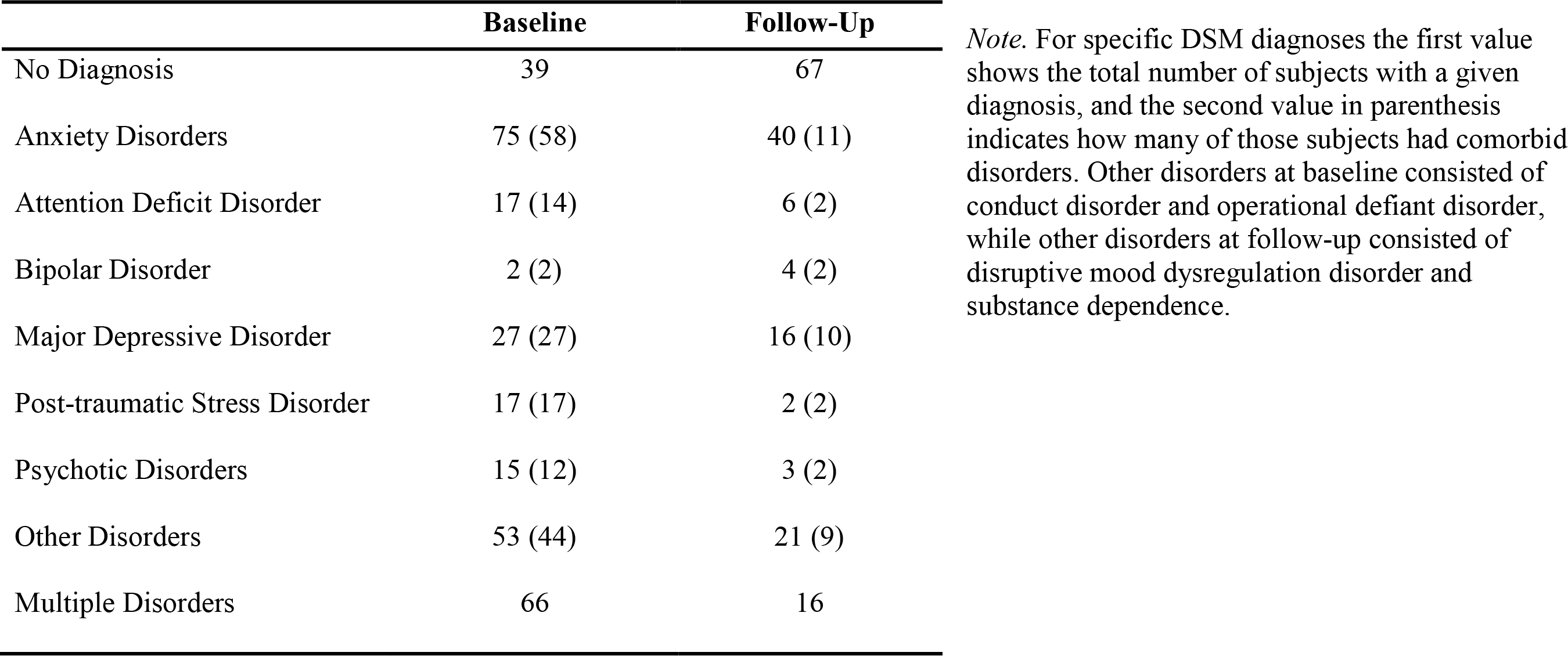
Number of participants with specific and comorbid disorders at baseline and follow-up.

**Table S2.** Sensitivity analyses of cross-sectional associations between structural covariance networks and irritability at follow-up.

| **NMF-Networks** | **Attention-Deficit**<br>**Hyperactivity** | **Anxiety**<br>**Symptoms** | **Depression**<br>**Severity** | **Excluding**<br>**Drug Users** |
| --- | --- | --- | --- | --- |
| N-1: Medial temporal lobe | .002** | <.001*** | <.001** | .003** |
| N-8: Temporo-parietal junction | .035 | .005* | .009* | .003** |
| N-9: Temporal pole & medial orbitofrontal cortex | .091 | .028* | .017* | .002** |
| N-10: Inferior temporal gyrus | .039 | .006** | .019* | .007** |
| N-11: Occipital cortex | .021* | .001** | .001** | .003** |
| N-13: Fusiform gyrus | <.001*** | <.001*** | <.001*** | <.001*** |
| N-15: Lateral temporal cortex | <.001** | .001** | <.001*** | <.001** |
| N-18: Lateral orbitofrontal cortex | .023* | .020* | .017* | .007** |
| N-20: Inferior temporal cortex | .123 | .018* | .027* | .006** |
| N-21: Lingual gyrus | .125 | .019* | .047* | .005** |
| N-22: Superior temporal gyrus | .111 | .012* | .017* | .009** |

*Note.* Uncorrected p-values are reported, and asterisks represent whether the associations survived FDR-correction: ( $p \leq .05$ )\*, ( $p \leq .01$ )\*\*, ( $p \leq .001$ )\*\*\*

**Table S3.** Sensitivity analyses of associations between longitudinal change within structural covariance networks and irritability at follow-up.

| **NMF-Networks** | **Attention-Deficit**<br>**Hyperactivity** | **Anxiety**<br>**Symptoms** | **Depression**<br>**Severity** | **Excluding**<br>**Drug Users** |
| --- | --- | --- | --- | --- |
| N-1: Medial temporal lobe | .001** | <.001*** | <.001*** | <.001*** |
| N-7: Inferior frontal cortex | .025* | .137 | .072 | .115 |
| N-9: Temporal pole and medial orbitofrontal cortex | .002** | .019* | .006* | .014* |
| N-11: Occipital cortex | <.001** | <.001*** | <.001*** | .004** |
| N-15: Lateral temporal cortex | .020* | .037 | .041 | .060 |
| N-17: Supramarginal gyrus | .041* | .065 | .058 | .111 |
| N-18: Lateral orbitofrontal cortex | .035* | .142 | .077 | .177 |
| N-20: Inferior temporal cortex | .021* | .018* | .015* | .006* |
| N-22: Superior temporal gyrus | .037* | .051 | .056 | .073 |

*Note.* Uncorrected p-values are reported, and asterisks represent whether the associations survived FDR-correction: ( $p \leq .05$ )\*, ( $p \leq .01$ )\*\*, ( $p \leq .001$ )\*\*\*

**Figure S1.**
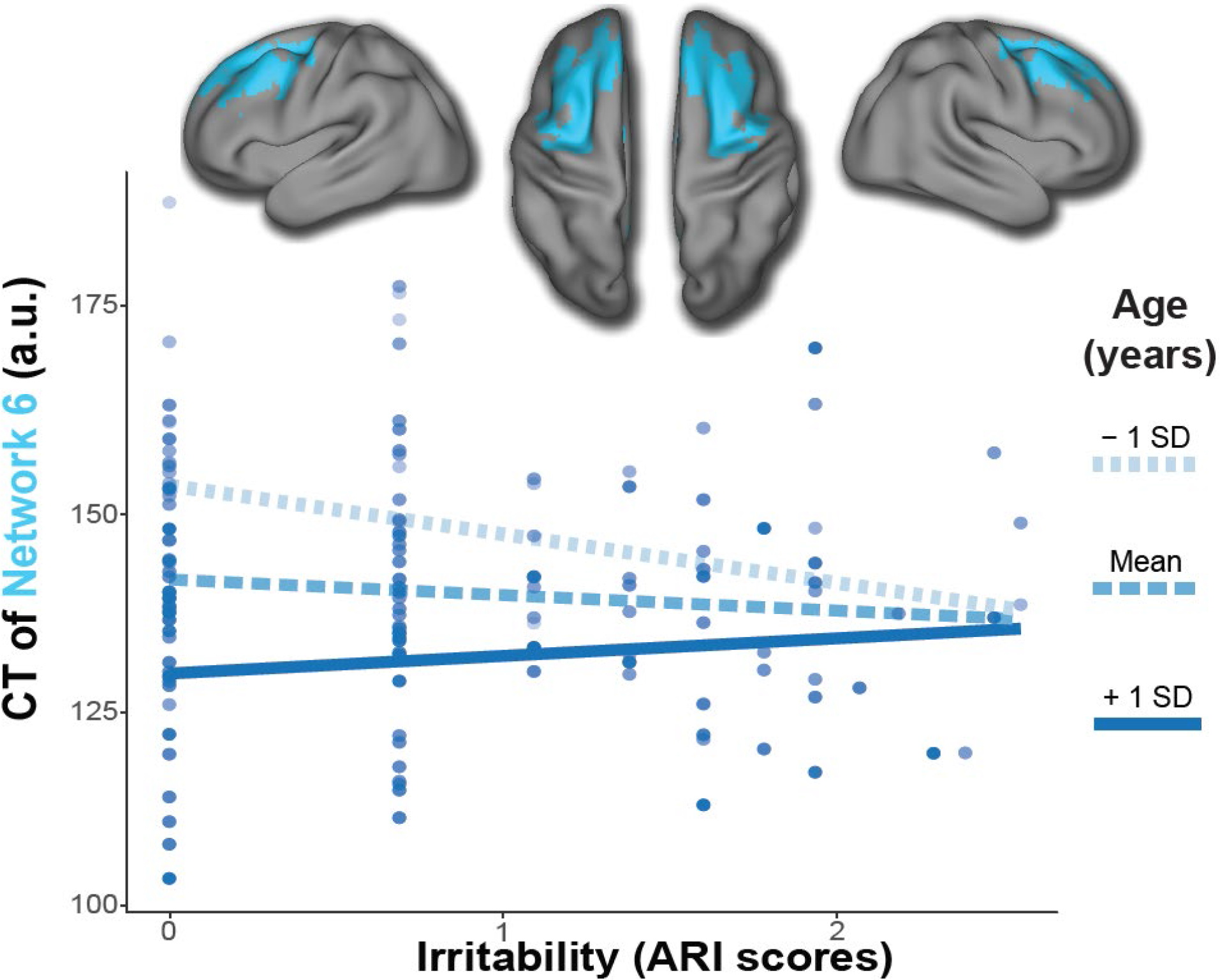
The relationship between irritability and the dorsolateral prefrontal cortex weakens with age. An irritability by age interaction effect significantly predicted CT of the covariance network encompassing the dorsolateral prefrontal cortex. A simple slopes analysis further revealed that the relationship between irritability and the thickness of the dorsolateral prefrontal cortex was significant among younger participants but not at the mean or +1 SD above the mean of age.

## ACKNOWLEDGEMENTS

This work was supported by grants from the National Institute of Mental Health: R01MH107703, R01MH113550, K99MH117274, R01MH112847, R01EB025471, R01MH113565, R01MH11207, R01MH11207, S10OD023495, and R01EB022573. Support for developing multivariate pattern analysis software was provided by a seed grant by the Center for Biomedical Computing and Image Analysis (CBICA) at Penn. Additional support was provided by the Lifespan Brain Institute at the Children’s Hospital of Philadelphia and Penn Medicine & the AE Foundation.

## REFERENCES

1. Leibenluft, E., Severe mood dysregulation, irritability, and the diagnostic boundaries of bipolar disorder in youths. Am J Psychiatry, 2011. 168(2): p. 129–42.

2. Stringaris, A., et al., Adult outcomes of youth irritability: a 20-year prospective community-based study. Am J Psychiatry, 2009. 166(9): p. 1048–54.

3. Stringaris, A., et al., Youth meeting symptom and impairment criteria for mania-like episodes lasting less than four days: an epidemiological enquiry. Journal of child psychology and psychiatry, and allied disciplines, 2010. 51(1): p. 31–38.

4. Biederman, J., et al., Further evidence of unique developmental phenotypic correlates of pediatric bipolar disorder: findings from a large sample of clinically referred preadolescent children assessed over the last 7 years. J Affect Disord, 2004. 82 Suppl 1: p. S45–58.

5. Stringaris, A., et al., Pediatric bipolar disorder versus severe mood dysregulation: risk for manic episodes on follow-up. J Am Acad Child Adolesc Psychiatry, 2010. 49(4): p. 397–405.

6. Leibenluft, E., et al., Chronic versus episodic irritability in youth: a community-based, longitudinal study of clinical and diagnostic associations. J Child Adolesc Psychopharmacol, 2006. 16(4): p. 456–66.

7. Leibenluft, E., et al., Defining clinical phenotypes of juvenile mania. Am J Psychiatry, 2003. 160(3): p. 430–7.

8. Research Domain Criteria (RDoC): Toward a New Classification Framework for Research on Mental Disorders. American Journal of Psychiatry, 2010. 167(7): p. 748–751.

9. Casey, B.J., M.E. Oliveri, and T. Insel, A Neurodevelopmental Perspective on the Research Domain Criteria (RDoC) Framework. Biological Psychiatry, 2014. 76(5): p. 350–353.

10. Brotman, M.A., et al., Prevalence, Clinical Correlates, and Longitudinal Course of Severe Mood Dysregulation in Children. Biological Psychiatry, 2006. 60(9): p. 991–997.

11. Dougherty, L.R., et al., Preschool irritability predicts child psychopathology, functional impairment, and service use at age nine. J Child Psychol Psychiatry, 2015. 56(9): p. 999–1007.

12. Belden, A.C., N.R. Thomson, and J.L. Luby, Temper tantrums in healthy versus depressed and disruptive preschoolers: defining tantrum behaviors associated with clinical problems. The Journal of pediatrics, 2008. 152(1): p. 117–122.

13. Keenan, K. and L.S. Wakschlag, More than the Terrible Twos: The Nature and Severity of Behavior Problems in Clinic-Referred Preschool Children. Journal of Abnormal Child Psychology, 2000. 28(1): p. 33–46.

14. S. Wakschlag, L., et al., Clinical Implications of a Dimensional Approach: The Normal:Abnormal Spectrum of Early Irritability. Vol. 54. 2015.

15. Pagliaccio, D., et al., Irritability Trajectories, Cortical Thickness, and Clinical Outcomes in a Sample Enriched for Preschool Depression. Journal of the American Academy of Child & Adolescent Psychiatry, 2018. 57(5): p. 336–342.e6.

16. Adleman, N.E., et al., Cross-sectional and longitudinal abnormalities in brain structure in children with severe mood dysregulation or bipolar disorder. J Child Psychol Psychiatry, 2012. 53(11): p. 1149–56.

17. Aron, A.R., et al., Converging evidence for a fronto-basal-ganglia network for inhibitory control of action and cognition. J Neurosci, 2007. 27(44): p. 11860–4.

18. Deveney, C.M., et al., Neural mechanisms of frustration in chronically irritable children. Am J Psychiatry, 2013. 170(10): p. 1186–94.

19. Tseng, W.-L., et al., Brain Mechanisms of Attention Orienting Following Frustration: Associations With Irritability and Age in Youths. American Journal of Psychiatry, 2018. 176(1): p. 67–76.

20. Wiggins, J.L., et al., Neural Correlates of Irritability in Disruptive Mood Dysregulation and Bipolar Disorders. American Journal of Psychiatry, 2016. 173(7): p. 722–730.

21. Deveney, C.M., et al., Neural recruitment during failed motor inhibition differentiates youths with bipolar disorder and severe mood dysregulation. Biological psychology, 2012. 89(1): p. 148–155.

22. Singh, M.K., et al., Neural correlates of response inhibition in pediatric bipolar disorder. J Child Adolesc Psychopharmacol, 2010. 20(1): p. 15–24.

23. Li, Y., et al., The neural substrates of cognitive flexibility are related to individual differences in preschool irritability: A fNIRS investigation. Dev Cogn Neurosci, 2017. 25: p. 138–144.

24. Perlman, S.B., et al., fNIRS evidence of prefrontal regulation of frustration in early childhood. Neuroimage, 2014. 85 Pt 1: p. 326–34.

25. Kringelbach, M.L., The human orbitofrontal cortex: linking reward to hedonic experience. Nature Reviews Neuroscience, 2005. 6: p. 691.

26. Goddard, G.V., Functions of the amygdala. Psychological Bulletin, 1964. 62(2): p. 89–109.

27. Fulwiler, C.E., J.A. King, and N. Zhang, Amygdala-orbitofrontal resting-state functional connectivity is associated with trait anger. Neuroreport, 2012. 23(10): p. 606–610.

28. Paus, T., M. Keshavan, and J.N. Giedd, Why do many psychiatric disorders emerge during adolescence? Nature reviews. Neuroscience, 2008. 9(12): p. 947–957.

29. Giedd, J.N., et al., Brain development during childhood and adolescence: a longitudinal MRI study. Nature Neuroscience, 1999. 2: p. 861.

30. Sotiras, A., S.M. Resnick, and C. Davatzikos, Finding imaging patterns of structural covariance via Non-Negative Matrix Factorization. Neuroimage, 2015. 108: p. 1–16.

31. Sotiras, A., et al., Patterns of coordinated cortical remodeling during adolescence and their associations with functional specialization and evolutionary expansion. Proc Natl Acad Sci U S A, 2017. 114(13): p. 3527–3532.

32. Satterthwaite, T.D., et al., The Philadelphia Neurodevelopmental Cohort: A publicly available resource for the study of normal and abnormal brain development in youth. NeuroImage, 2016. 124: p. 1115–1119.

33. Satterthwaite, T.D., et al., Neuroimaging of the Philadelphia neurodevelopmental cohort. Neuroimage, 2014. 86: p. 544–53.

34. Calkins, M.E., et al., The psychosis spectrum in a young U.S. community sample: findings from the Philadelphia Neurodevelopmental Cohort. World Psychiatry, 2014. 13(3): p. 296–305.

35. Kaufman, J., et al., Schedule for Affective Disorders and Schizophrenia for School-Age Children-Present and Lifetime Version (K-SADS-PL): initial reliability and validity data. J Am Acad Child Adolesc Psychiatry, 1997. 36(7): p. 980–8.

36. Calkins, M.E., et al., The Philadelphia Neurodevelopmental Cohort: constructing a deep phenotyping collaborative. J Child Psychol Psychiatry, 2015. 56(12): p. 1356–1369.

37. Calkins, M.E., et al., Persistence of psychosis spectrum symptoms in the Philadelphia Neurodevelopmental Cohort: a prospective two-year follow-up. World psychiatry: official journal of the World Psychiatric Association (WPA), 2017. 16(1): p. 62–76.

38. First, M., et al., Structured clinical interview for DSM-IV-TR Axis I Disorders, Research Version, Non-patient Edition. 2002.

39. Stringaris, A., et al., The Affective Reactivity Index: a concise irritability scale for clinical and research settings. J Child Psychol Psychiatry, 2012. 53(11): p. 1109–17.

40. Mulraney, M.A., G.A. Melvin, and B.J. Tonge, Psychometric properties of the affective reactivity index in Australian adults and adolescents. Psychol Assess, 2014. 26(1): p. 148–55.

41. Beck, A.T., R.A. Steer, and G.K. Brown, Beck depression inventory-II. San Antonio, 1996. 78(2): p. 490–498.

42. Birmaher, B., et al., The screen for child anxiety related emotional disorders (SCARED): Scale construction and psychometric characteristics. Journal of the American Academy of Child & Adolescent Psychiatry, 1997. 36(4): p. 545–553.

43. Swanson, J.M., et al., Categorical and Dimensional Definitions and Evaluations of Symptoms of ADHD: History of the SNAP and the SWAN Rating Scales. The International journal of educational and psychological assessment, 2012. 10(1): p. 51.

44. Tustison, N.J., et al., Large-scale evaluation of ANTs and FreeSurfer cortical thickness measurements. Neuroimage, 2014. 99: p. 166–79.

45. Ciric, R., et al., Benchmarking of participant-level confound regression strategies for the control of motion artifact in studies of functional connectivity. NeuroImage, 2017. 154: p. 174–187.

46. Tustison, N.J., et al., N4ITK: improved N3 bias correction. IEEE Trans Med Imaging, 2010.29(6): p. 1310–20.

47. Avants, B.B., et al., A reproducible evaluation of ANTs similarity metric performance in brain image registration. NeuroImage, 2011. 54(3): p. 2033–2044.

48. Klein, A., et al., Evaluation of volume-based and surface-based brain image registration methods. Neuroimage, 2010. 51(1): p. 214–20.

49. Das, S.R., et al., Registration based cortical thickness measurement. Neuroimage, 2009.45(3): p. 867–79.

50. Rosen, A.F.G., et al., Quantitative assessment of structural image quality. NeuroImage, 2018. 169: p. 407–418.

51. Alexander-Bloch, A., J.N. Giedd, and E. Bullmore, Imaging structural co-variance between human brain regions. Nat Rev Neurosci, 2013. 14(5): p. 322–36.

52. Zielinski, B.A., et al., Network-level structural covariance in the developing brain. Proceedings of the National Academy of Sciences, 2010. 107(42): p. 18191–18196.

53. Eklund, A., T.E. Nichols, and H. Knutsson, Cluster failure: Why fMRI inferences for spatial extent have inflated false-positive rates. Proceedings of the National Academy of Sciences, 2016. 113(28): p. 7900–7905.

54. Yang, Z. and E. Oja, Linear and Nonlinear Projective Nonnegative Matrix Factorization. IEEE Transactions on Neural Networks, 2010. 21(5): p. 734–749.

55. Boutsidis, C. and E. Gallopoulos, Gallopoulos, E.: Svd based initialization: A head start for nonnegative matrix factorization. Pattern Recognition 41(4), 1350-1362. Vol. 1350-1362. 2008. 1350–1362.

56. Van Essen, D.C., et al., An integrated software suite for surface-based analyses of cerebral cortex. J Am Med Inform Assoc, 2001. 8(5): p. 443–59.

57. Johnson, W.E., C. Li, and A. Rabinovic, Adjusting batch effects in microarray expression data using empirical Bayes methods. Biostatistics, 2007. 8(1): p. 118–127.

58. Fortin, J.-P., et al., Harmonization of cortical thickness measurements across scanners and sites. NeuroImage, 2018. 167: p. 104–120.

59. Fortin, J.-P., et al., Harmonization of multi-site diffusion tensor imaging data. NeuroImage, 2017. 161: p. 149–170.

60. Yu, M., et al., Statistical harmonization corrects site effects in functional connectivity measurements from multi-site fMRI data. Hum Brain Mapp, 2018. 39(11): p. 4213–4227.

61. R Development Core Team, R: A language and environment for statistical computing. 2018: R Foundation for Statistical Computing. Retrieved from http://www.R-project.org.

62. Lenroot, R.K., et al., Sexual dimorphism of brain developmental trajectories during childhood and adolescence. Neuroimage, 2007. 36(4): p. 1065–73.

63. Wood, S.N. and N.H. Augustin, GAMs with integrated model selection using penalized regression splines and applications to environmental modelling. Ecological modelling, 2002. 157(2-3): p. 157–177.

64. Cannon, T.D., et al., Progressive reduction in cortical thickness as psychosis develops: a multisite longitudinal neuroimaging study of youth at elevated clinical risk. Biological psychiatry, 2015. 77(2): p. 147–157.

65. Winston, J.S., J. O’Doherty, and R.J. Dolan, Common and distinct neural responses during direct and incidental processing of multiple facial emotions. NeuroImage, 2003. 20(1): p. 84–97.

66. Brotman, M.A., et al., Amygdala activation during emotion processing of neutral faces in children with severe mood dysregulation versus ADHD or bipolar disorder. The American journal of psychiatry, 2010. 167(1): p. 61–69.

67. Guyer, A.E., et al., Specificity of facial expression labeling deficits in childhood psychopathology. J Child Psychol Psychiatry, 2007. 48(9): p. 863–71.

68. Kircanski, K., et al., A Latent Variable Approach to Differentiating Neural Mechanisms of Irritability and Anxiety in Youth. JAMA Psychiatry, 2018. 75(6): p. 631–639.

69. Gold, A.L., et al., Comparing Brain Morphometry Across Multiple Childhood Psychiatric Disorders. J Am Acad Child Adolesc Psychiatry, 2016. 55(12): p. 1027–1037.e3.

70. Mueller, K., et al., Commentary: Cluster failure: Why fMRI inferences for spatial extent have inflated false-positive rates. Frontiers in human neuroscience, 2017. 11: p. 345–345.

